# Bayesian inference of chromatin structure ensembles from population Hi-C data

**DOI:** 10.1101/493676

**Authors:** Simeon Carstens, Michael Nilges, Michael Habeck

**Affiliations:** Statistical Inverse Problems in Biophysics, Max Planck Institute for Biophysical Chemistry, Am Faßberg 11, 37077 Göottingen; Department of NMR based Structural Biology, Max Planck Institute for Biophysical Chemistry, Am Faßberg 11, 37077 Göottingen; Unit´e de Bioinformatique Structurale, Institut Pasteur, Rue du Dr. Roux 25–28, 75015 Paris; Felix Bernstein Institute for Mathematical Statistics in the Biosciences, University of Göottingen, Göottingen, Germany

## Abstract

High-throughput chromosome conformation capture (Hi-C) experiments are typically performed on a large population of cells and therefore only yield average numbers of genomic contacts. Nevertheless population Hi-C data are often interpreted in terms of a single genomic structure, which ignores all cell-to-cell variability. We propose a probabilistic, statistically rigorous method to infer chromatin structure ensembles from population Hi-C data that takes the ensemble nature of the data explicitly into account and allows us to infer the number of structures required to explain the data.

---

In recent years, chromosome conformation capture (3C) methods have emerged as a powerful tool to investigate the three-dimensional organization of genomes on previously inaccessible length scales. Thanks to experimental advances and decreasing sequencing costs, genome-wide 3C methods such as Hi-C (1) and Hi-C variants (2; 3) have yielded contact maps with resolutions of up to 1 kb. High-resolution contact maps have provided fascinating insights into the organization of chromatin domains and compartments of various sizes and their role in gene regulation (1; 4; 5; 6; 3).

Although Hi-C can be performed in single cells (7; 8; 9; 10; 11), much richer data are usually obtained by analyzing populations of cells at the price of a more difficult interpretation of the data: Contact maps obtained in population Hi-C experiments show an average over millions of cells. Given that single-cell Hi-C experiments revealed significant structural variability among cell nuclei (7), it remains difficult to assess the information content and degree of structural heterogeneity reflected by population Hi-C data.

Much effort has gone into the development of chromatin structure determination approaches, in which a single consensus structure is calculated (12). In light of recent results on cell-to-cell variability and considering experimental limitations and biases, consensus structure approaches are fundamentally limited, if not flawed (13). Several approaches taking into account the heterogeneity of cell populations have been explored. Matrix deconvolution has been used to unmix Hi-C matrices without explicitly modeling 3D structures (14; 15). While some structural information is already apparent in contact matrices, it is useful to find an approximate 3D representation giving rise to the data. For example, a 3D chromosome model allows the detection of three-way interactions not accessible in 3C data (16). Furthermore, data from other sources (e.g., imaging) can be included in the modeling process. A second approach is thus to calculate an ensemble of structures that reflect the structural heterogeneity of chromosomes more realistically than consensus structure approaches and reproduce the contact map only on average. This can be achieved in two different ways: First, one can optimize a set of structures generated from a polymer model so as to reproduce, on average, the experimental data (2; 17; 18; 19). A second possibility is to adjust the parameters of a polymer model such that by simulating the polymer model an ensemble of structures is obtained that reproduces the experimental data on average (20; 21; 22).

In this communication, we extend the scope of our previous work on Bayesian chromosome structure inference from single-cell Hi-C data (23) to population-averaged contact data, combining the explicit modeling of chromatin structure populations with a rigorous probabilistic interpretation of the data. We leverage the full power of Bayesian inference to compute multi-state models of chromosome structures, which together reproduce the population contact data.

Our approach is based on the Inferential Structure Determination (ISD) (24, *Supplementary Notes*) framework. ISD infers a structural model *x* along with unknown modeling parameters *α* from experimental data *D* and background information *I*. The inference problem is solved by sampling from the joint posterior distribution for *x* and *α*, conditioned on *D* and *I*, which can be expressed using Bayes’ theorem as

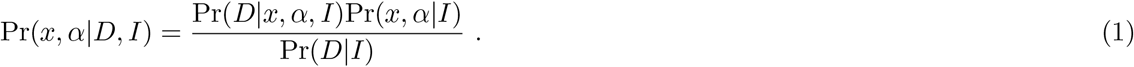

Here we assume that a single structure *x* can explain the data *D*, as is the case for, e.g., proteins with a stable and unique fold. Sampling from Pr(*x, α|D, I*) results in a statistically well-defined ensemble of structures with associated estimates of nuisance parameters *α*, each being assigned a probability given by the posterior distribution.

To accommodate the fact that most 3C experiments are performed on populations of molecules with possibly very different conformations due to cell-to-cell heterogeneity and dynamics, we extend the scope of the ISD approach to the inference of multi-state models *X* = (*x*^1^, …, *x*^*n*^) consisting of *n* structures *x*^*k*^. The central quantity in this extended ISD framework is the posterior distribution

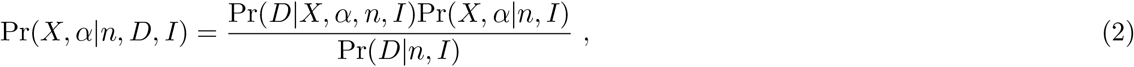

sampling from which results in a *hyperensemble*, that is, a representative set of multi-state models *X*_*i*_ with associated nuisance parameters *α*_*i*_, in which each pair (*X*_*i*_, *α*_*i*_) has a probability given by Equation 2. In the case of a single state (*n* = 1), we recover the original ISD formulation and infer a consensus structure.

At the center of a Bayesian approach is a generative model for the data Pr(*D|X, α, n, I*) given a set *X* of *n* structures. Our generative model is based on a forward model to compute average contact matrices, which are idealized, noise-free data. As shown in Figure 1a, each structure *x*^*k*^ that is part of *X* gives rise to a smoothed contact matrix based on a threshold distance. The *n* contact matrices are summed together and multiplied with a scale factor *α* that is estimated from the data. We model deviations from these idealized data due to experimental noise, forward model inaccuracy and, in case of Hi-C data, data normalization with Poisson distributions whose rates are given by the idealized count data.

**Figure 1:**
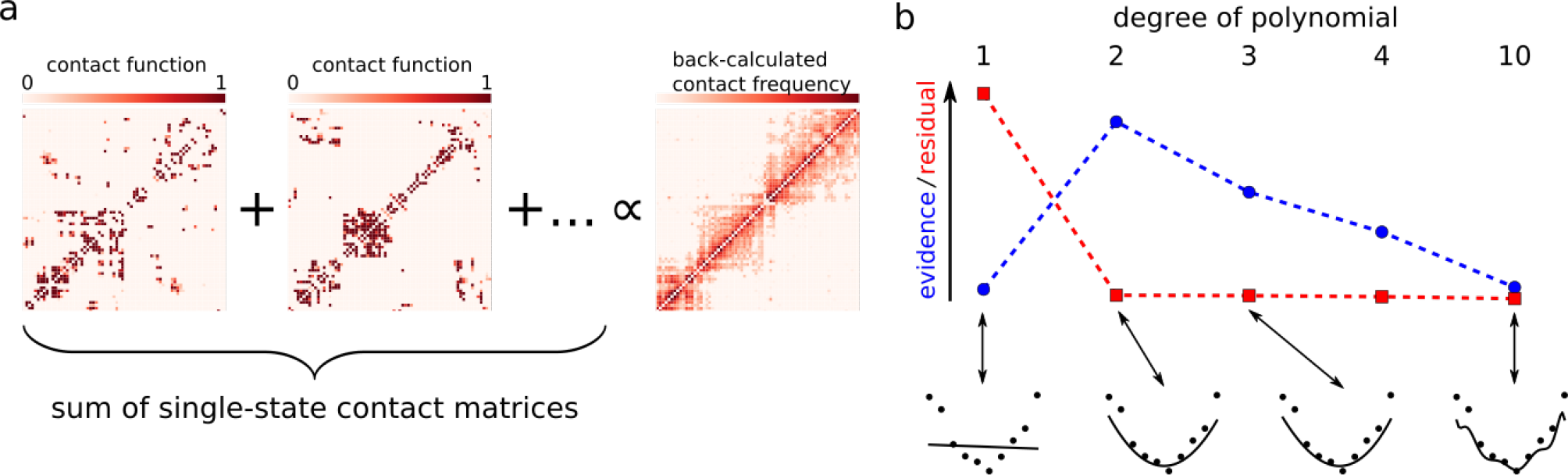
Main components of the proposed chromatin structure inference approach. **(a)**Forward model to calculate idealized contact frequency matrices from structures that are part of a multi-state model. The contact matrices arising from each structure are summed together and multiplied with a scaling factor to yield the backcalculated contact frequency matrix on the right. **(b)**Bayesian model comparison illustrated for a polynomial curve fitting example. Shown is the evidence (blue) and the residual mismatch between model and data (red) for polynomials of increasing degree. Data points and best fitting models are shown below the *x* axis.

Under reasonable independence assumptions, the joint prior distribution Pr(*X, α|n, I*) factorizes and we can formulate separate prior distributions for all states (*x*^1^, …, *x*^*n*^) and for the scale factor *α*. We model the physical interactions within a single structure *x*^*k*^ by a coarse-grained beads-on-a-string model similar to our model for single-cell data (23). Further details on our statistical model for population-averaged contact data and our choice of prior distributions are explained in *Supplementary Notes*.

We use a range of Markov Chain Monte Carlo (MCMC) techniques to address the challenging task of sampling from Pr(*X, α|n, D, I*) (see *Supplementary Notes* for details). Once representative samples have been drawn from the posterior distribution, the structure determination problem is solved and the estimated multi-state models can be analyzed.

A crucial parameter in all structure calculation methods that model explicit ensembles (2; 17; 18) is the number of states *n*. Existing multi-state modeling approaches choose *n* in an *ad hoc* manner. For example, in the GEM (18) software, the default value for *n* is four and differs largely from *n* = 10000 used by Alber and coworkers (17). From a Bayesian perspective, the number of states *n* is a model parameter that should be inferred from the data. Depending on the true heterogeneity of the chromosome structures *n* will adopt smaller or larger values. If *n* is too small, we underfit the contact data resulting in highly strained structural models. If *n* is too large, the structures of the states are only loosely defined and potentially overfit the contact data. Therefore, a simple goodness-of-fit criterion is not sufficient to select *n*. We have to balance goodness-of-fit against model complexity. Figure 1b illustrates this point for polynomial curve fitting. The degree of the polynomial determines the flexibility of the fitting function and therefore corresponds to the number of states *n*. For data generated from a parabola, the goodness-of-fit reaches an optimum if we use polynomials of degree two and higher eventually resulting in overfitting: the fitted curve closely follows each data point.

A unique feature of Bayesian data analysis is that it offers an objective, albeit computationally often challenging framework for model comparison (25). Our belief in a particular choice of *n* is encoded by the posterior probability

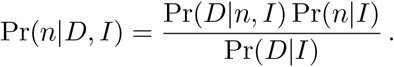

Assuming that *a priori* all *n* are equally likely, the ratio of two choices for the number of states *n* and *n*′ can be written as

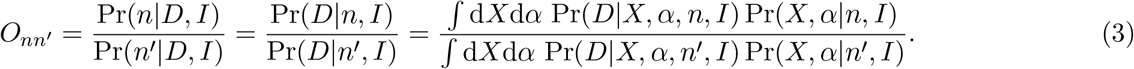

The odds ratio *O*_*nn′*_ is called the *Bayes factor* and tells us which model to prefer: for *O*_*nn′*_ > 1, an *n*-state model describes the data better; while *O*_*nn′*_ < 1 indicates a preference for a model based on *n′* states. The model evidence Pr(*D|n, I*) has a built-in penalty for model complexity. In the curve fitting example (Figure 1b), Pr(*D|n, I*) selects the correct degree of the polynomial (*n* = 2), although polynomials of degree > 2 achieve a better fit. This is because with increasing degree also the number of curves that do not fit the data increases, which is reflected by a decrease in the model evidence. The evaluation of the model evidence involves calculating two possibly intractable and high-dimensional integrals, which in all but the most trivial cases have to be approximated numerically. As several of the occurring distributions are of a non-standard form, we use histogram reweighting (26; 27) to approximate the evidences.

We implemented our method in a freely available Python package (http://bitbucket.org/simeon_carstens/ensemble_hic). As a proof of concept, we first apply our inference approach to contact data simulated from two protein structures. We consider beads-on-a-string models of two protein domains, the B1 immunoglobin-binding domain of streptococcal protein G (PDB identifier 1PGA) and the SH3 domain of human Fyn (PDB identifier 1SHF). Only positions of *C*_*α*_ atoms were taken into account, and the SH3 domain was truncated by three C-terminal residues to match the length of 1PGA. This results in two very different conformations of a chain of 56 beads (Figure 2a,b). We generated simulated contact frequencies and used ISD to infer representative multi-state models. We simulated the posterior distributions for *n* = 1, 2, 3, 4, 5, 10 states. Model comparison shows that the model with *n* = 2 states is strongly preferred, as indicated by *P* (*D|n* = 2, *I*)/*P* (*D|n* = 1,*I*) > 10^307^ and *P* (*D|n* = 2,*I*)/*P* (*D|n* = 3,*I*) > 10^160^ (Figure 2c). It is reassuring that while the evidence decreases for *n* > 2, models with an even number of states are preferred over models with a similar, but uneven number of states. The reason for this is that for even *n*, half of the states adopt the conformation of the *B*1 domain and the other half adopts the conformation of the SH3 domain, thus having few superfluous parameters (Supplementary Figure S1). For uneven *n*, on the other hand, there is at least one superfluous state (Supplementary Figures S2 and S3), contributing uninformative parameters which are penalized in the evidence. We find that, for *n* > 2, the average scale factor *α* decreases monotonically with an increasing number of states (Figure 2f), in agreement with our intuition that *α* compensates for the smaller number of simulated states as compared to typical numbers of counts in the data.

**Figure 2:**
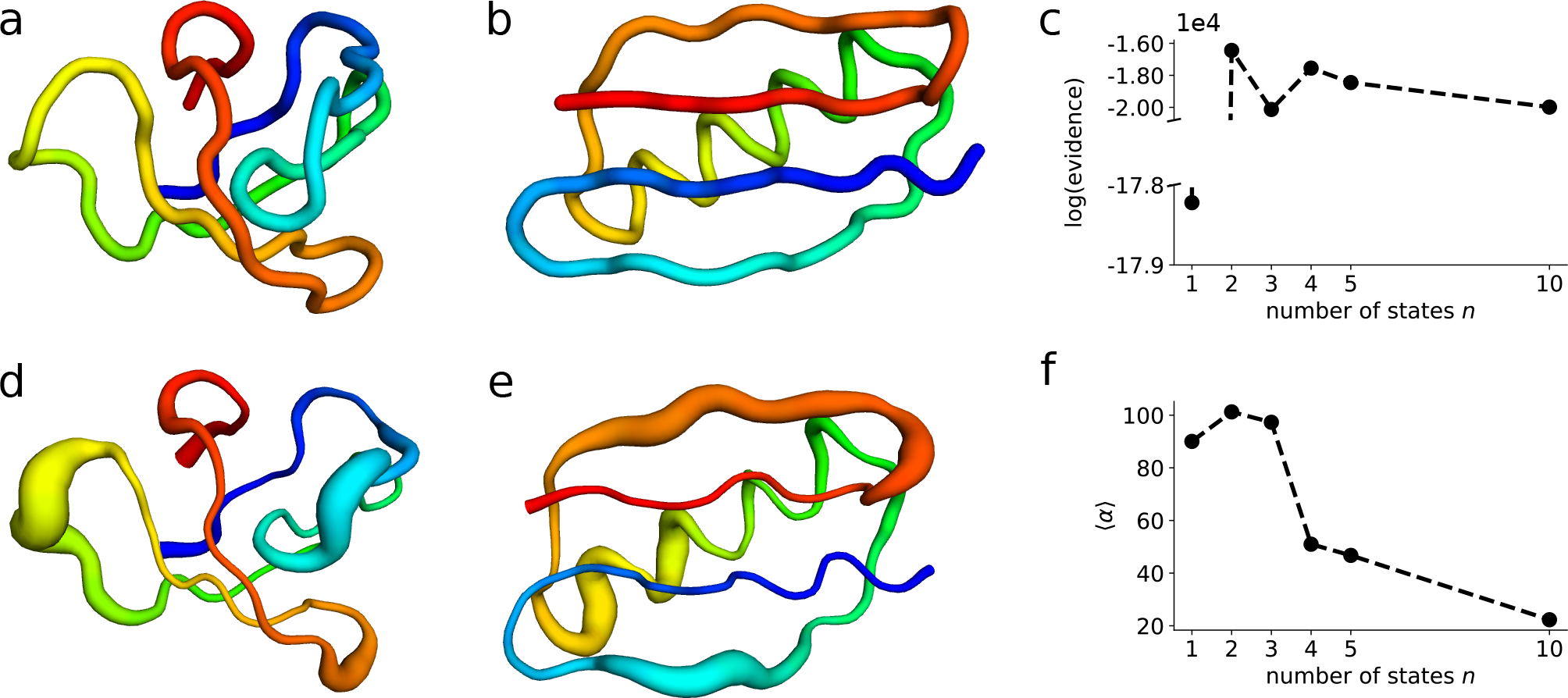
Reconstruction of protein structures from simulated data. **(a), (b)**Reference structures of the SH3 domain **(a)** and protein G **(b)**. **(c)** Model evidences for different number of states. **(d), (e)** “Sausage representation” of reconstructed folds (*n* = 2 states). Tube thickness indicates the local precision of bead positions. **(f)**Average values of sampled scale factor.

Figure 2d,e shows the two reconstructed states from the *n* = 2 simulation which deviate from the ground truth by an root-mean-square deviation of bead positions (RMSD) of (1.46 ± 0.11) Å for the B1 domain and (1.59 ± 0.21) Å for the SH3 domain. ISD thus recovers both conformations to very good accuracy.

Having established the validity of our approach on simulated data, we now apply our extended ISD approach to a 5C dataset covering the X-inactivation center in mouse embryonic stem cells (mESCs)(5). We restrict our analysis to contact frequencies between primer sites within the two consecutive TADs harbouring the *Tsix* and *Xist* promoters, respectively, and their boundary region. Similar to previous work on the same data (20), we model the corresponding ~920 kb region by *M* beads, each representing a genomic length of ~ 920/M kb. To keep the computational efforts for sufficient MCMC sampling reasonable, we chose a rather coarse resolution with one bead representing 15 kb, resulting in a total of 62 beads. Simulating the ISD posterior distribution for *n* = 1, 5, 10, 20, 30, 40, and 100 states and calculating model evidences, we find that the multi-state model with *n* = 20 states is clearly preferred by the data (Figure 3a). This result establishes that Bayesian model comparison is indeed able to determine a unique, optimal number of states for 5C data. The low resolution of these simulations lets us expect limited biological meaning of the resulting structures. To prove that a larger number of beads and thus higher resolution yields biologically meaningful structures, we increased the number of beads to *M* = 308. Our model for the chromatin polymer then coincides with the model chosen in previous work (20) on the same dataset. In the following, we thus analyze high-resolution chromatin structure ensembles obtained from a posterior simulation for *n* = 20. Out of three simulations with identical parameters (but different, random initial state), we chose the one with the highest mean posterior probability. While the optimal number of states for low- and high-resolution models does not neccessarily have to match, for *n* = 20, the back-calculated data reproduces the experimental data very well, with most mismatches stemming from short-range contacts contributing little structural information (Supplementary Figure S4). Determining the optimal number of states at the 3 kb resolution is currently not feasible, because accurate and exhaustive sampling of very high-dimensional posterior distributions is required for calculating good estimates of the evidence. This is outside the scope of our current implementation and this contribution.

**Figure 3:**
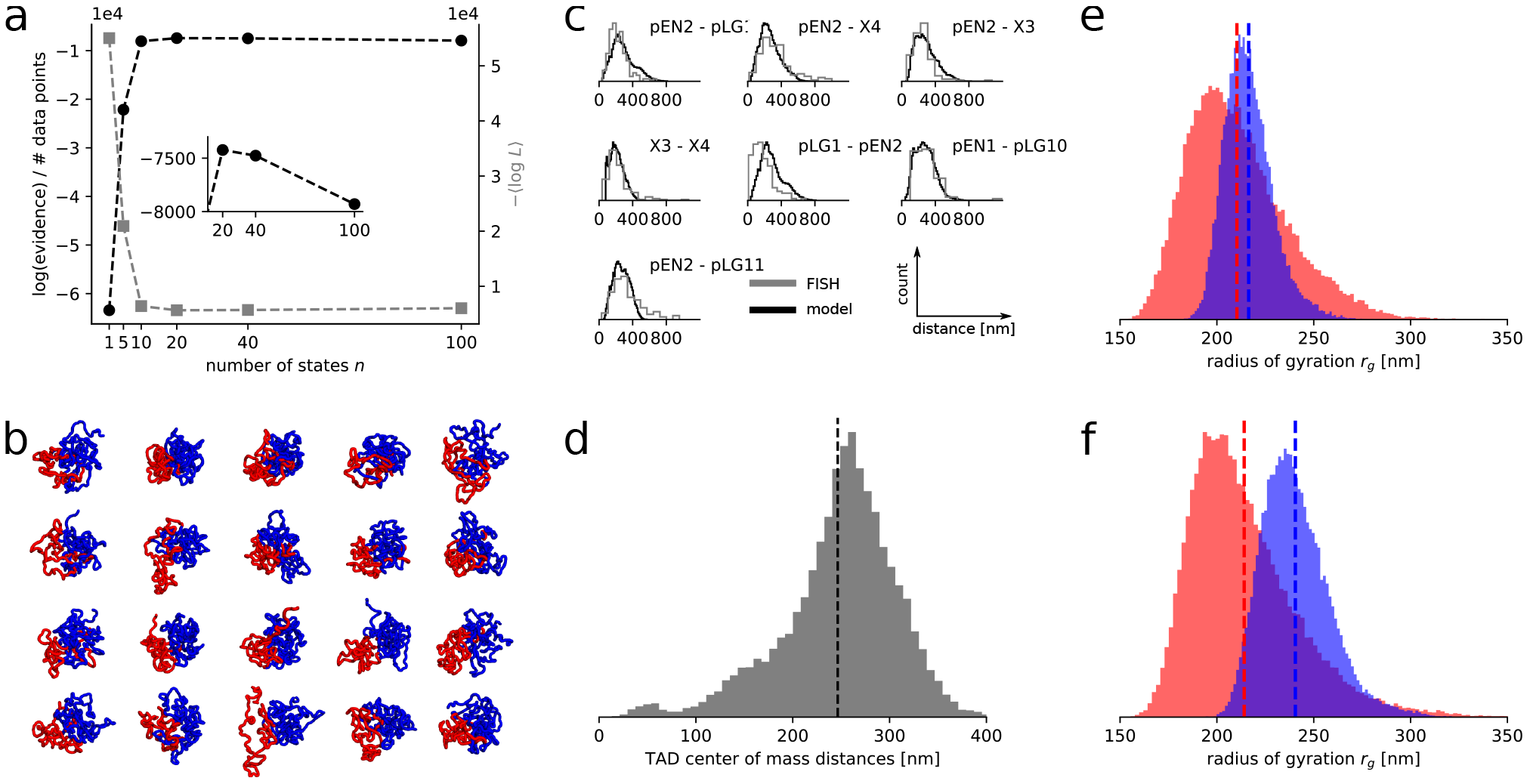
Inference of multi-state chromatin models from 5C data on male **(a–d)**and female **(e, f)** mESCs. **(a)** Model evidences and average likelihoods for various number of states at 15 kb resolution. Data in all other panels of this figure are calculated from a *n* = 20 simulation at 3 kb resolution. **(b)** Representative sample. Here and in the following panels, results for *Tsix* TAD in red and for *Xist* TAD in blue. **(c)** Pairwise distances between several loci as measured by DNA FISH and in inferred multi-state models. Names of genes overlapping the probes are shown. **(d)** Distance between centers of mass of *Tsix* and *Xist* TADs. The dashed line indicates the average radius of gyration of the whole genomic region. **(e, f)** Distribution of the radius of gyration for data obtained before **(e)** onset of differentiation and 48h later **(f)**. Dashed lines indicate sample means

Visual inspection of sample populations (Figure 3b) reveals a large structural variability which cannot possibly be captured by only a few, save one single structure. This heterogeneity is already apparent in the low-resolution models (Supplementary Figure S5). Before further analyzing our models, we validate the inferred multi-state chromatin models with previously measured FISH data (20). Figure 3c shows that experimental distances are in excellent agreement with distances obtained from the simulated structures, confirming that our approach yields structures matching data from experimental cell populations not only on average, but also correctly reproducing the spread of the population. Given the conformational heterogeneity in our structures, an interesting question our multi-state model approach allows us to answer is how homogeneous the compartmentalization of the genome into TADs is across a cell population. Figure 3d indicates that the *Tsix* and *Xist* TADs often intermingle considerably. Further validation of our modeling approach comes from considering data obtained from female mESCs before and 48h after onset of differentiation (5). In this cell line, transcription of the *Xist* TAD is upregulated while transcription levels of the Tsix TAD remain the same (5). After again choosing *n* = 20 and simulating the corresponding posterior distribution, measurement of the radii of gyration of both TADs reveals that before differentiation (Figure 3e), the *Xist* TAD assumes more densely packed conformations as opposed to after differentiation (Figure 3f), while the packing of the *Tsix* TAD stays almost the same. This agrees with the notion that in order to be highly transcribed, chromatin has to be loosely packed so as to provide easy access for the transcription machinery.

Finally, we compare the structures estimated by our extended ISD approach to the result of two previous approaches, which also model 3C data by multi-state models. We modified the publicly available code for PGS (28), the software used by Alber and co-workers (17), so as to work with the genomic region considered in this work. Calculations for three different numbers of states show good agreement of the resulting structure populations with afore-mentioned FISH data (20) for some loci pairs, while for others, the distance distributions differ significantly (Supplementary Figure S6). On average, the structure populations calculated with PGS show similar radii of gyration as our structures, but the radius of gyration distributions of the PGS structure populations is much broader. It is noteworthy that for structure populations calculated using PGS, neither FISH loci distance nor radius of gyration histograms vary significantly with the number of states, which span two orders of magnitude. Distances between most FISH loci are significantly different in both ISD and PGS structure populations as compared to a simulation of the null model without any data given by the polymer model used in our ISD approach (Supplementary Figure S6). We also calculated structure populations using GEM (18), but found that, irrespective of the number of states, which we varied between 4 and 100, there is virtually no heterogeneity in the populations (data not shown).

In conclusion, using both simulated and 5C data, we showed that the proposed Bayesian approach to chromatin structure inference is able to construct multi-state models from contact data that agree well with independent biological observations. Our method comes with an objective measure for the number of states required to accurately model population-based data, a feature previous approaches (2; 17; 18) are missing. The number of states used in previous work varies from very low numbers of states (*n* = 4) (18) and very large populations (*n* = 10^4^) (2; 17). Bayesian model comparison can, provided sufficiently accurate MCMC sampling, uniquely and objectively determine this number. While our method includes consensus structure calculation as a special case, it clearly demonstrates that this kind of data is much more appropriately modeled using a larger number of structures. At the same time, the Bayesian model comparison approach employed in this work helps to avoid overfitting the data.

This work opens up several avenues of further research. A more efficient implementation using toolkits such as OpenMM (29), possibly leveraging the power of graphics processing units (GPUs), would increase the feasible range of both resolutions and system size. More detailed structural prior information encoding, e.g., *a priori* contact probabilities between loci based on epigenetic marks (30; 31) and more detailed forward- and error models for the data, possibly including Hi-C data normalization routines, would likely result in more accurate 3D models. Eventually, with these improvements, the model comparison approach implemented in this work could rise from merely helping to set an important modeling parameter to a measure of cell-to-cell variability with the potential to answer a plethora of important biological questions. Finally, we note that our approach is not limited to 3C data, but could also be useful for other kinds of contact data in which structural heterogeneity can be suspected, for example crosslinking studies of protein complexes.

## Acknowledgements

We thank Luca Giorgetti for sharing DNA-FISH data. SC and MN acknowledge funding from the Agence National de la Recherche (ANR-11-MONU-0020). SC thanks Christian Griesinger for funding and support. MH acknowledges funding by the German Research Foundation (DFG) under grant SFB 860, TP B09.

## Supplementary Information

### Supplementary Figures

**Figure S1:**
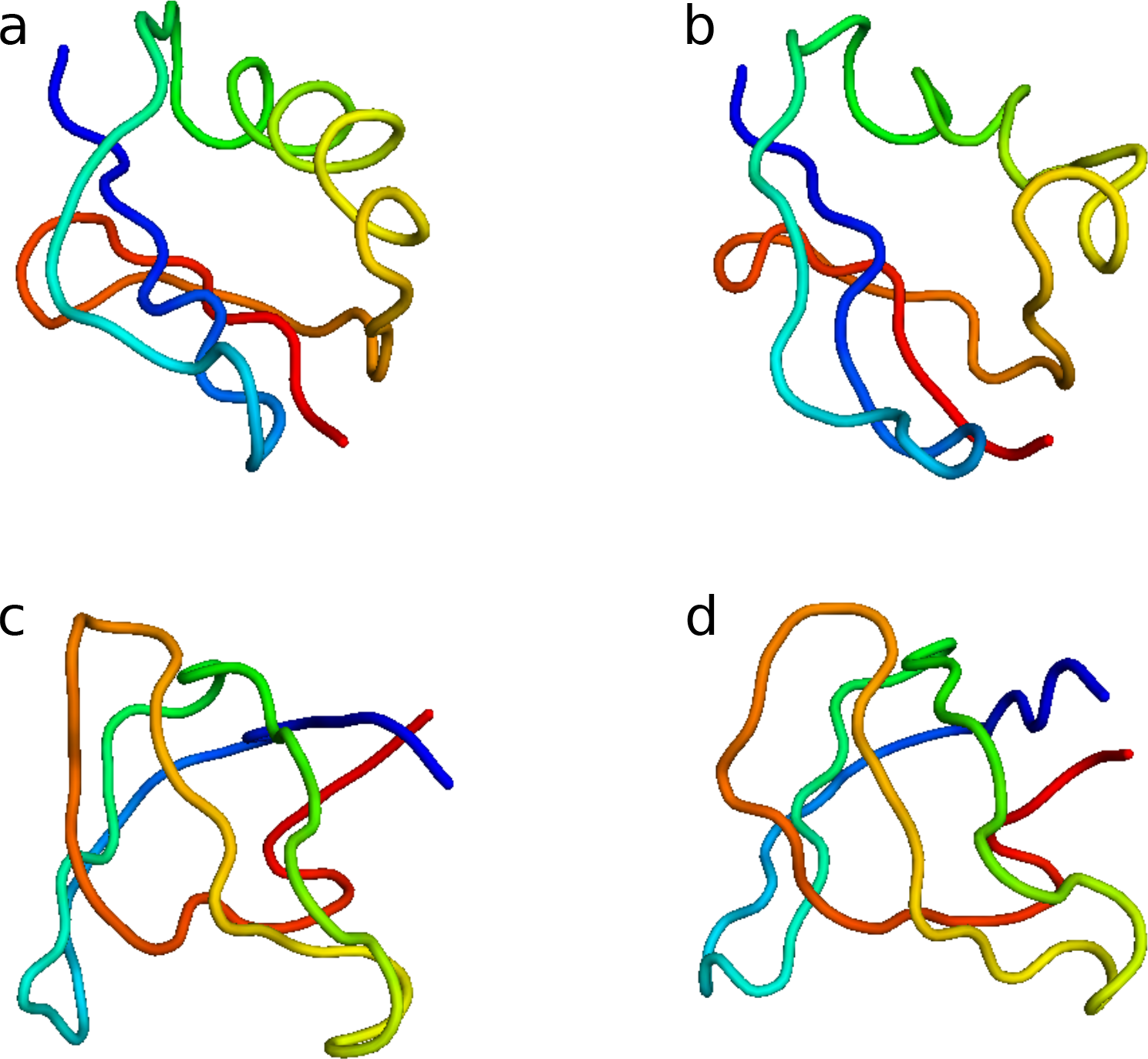
Reconstruction of protein structures from simulated contact frequency data. Shown is the Maximum *a posteriori* (MAP) estimate for *n* = 4 states. **(a,b)**: states approximating the conformation of 1PGA. **(c,d)**: states assuming the conformation of 1SHF.

**Figure S2:**
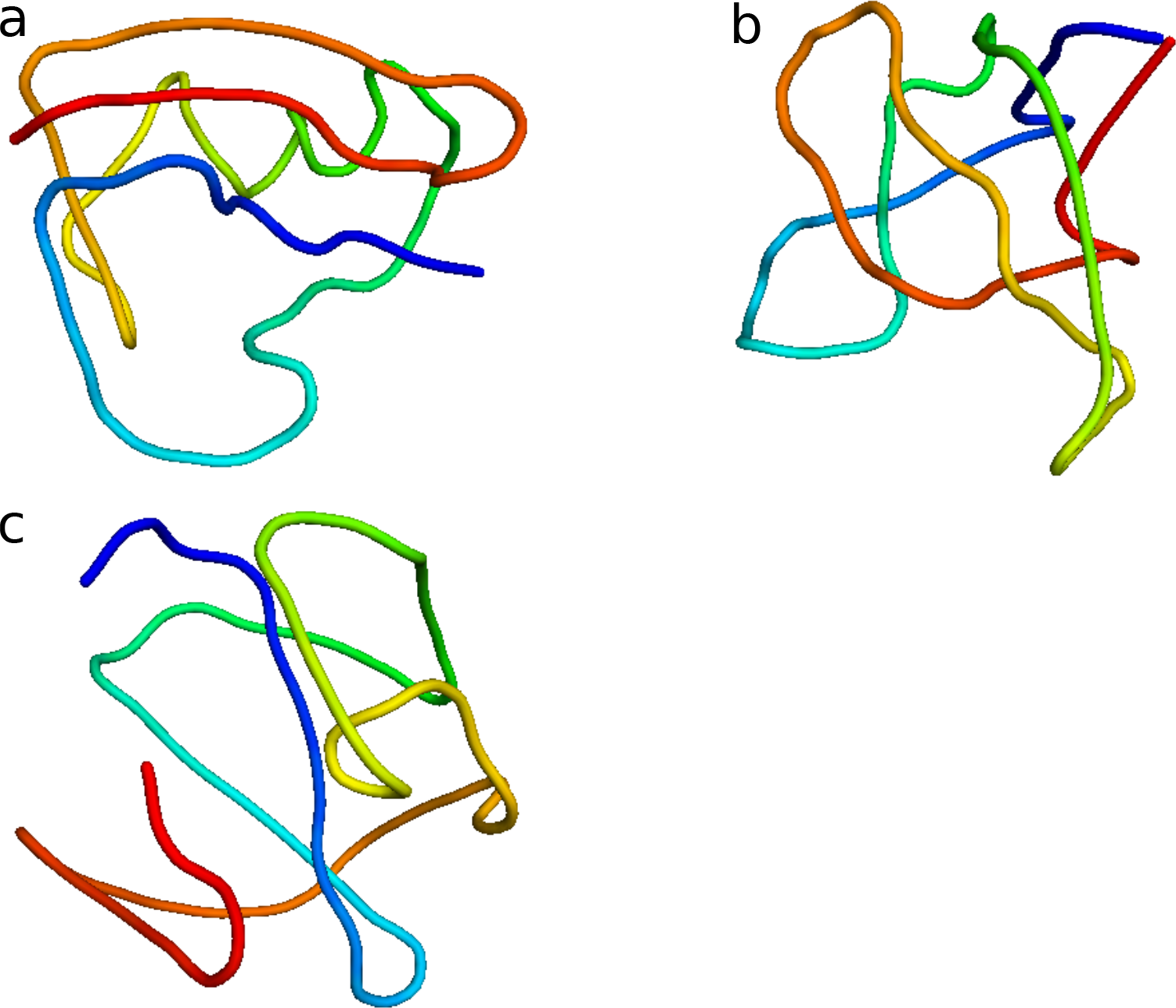
Reconstruction of protein structures from simulated contact frequency data. Shown is the MAP estimate for *n* = 3 states. **(a)**: state approximating the conformation of 1PGA. **(b)**: state approximating the conformation of 1SHF. **(c)**: “superfluous”, uninformative state.

**Figure S3:**
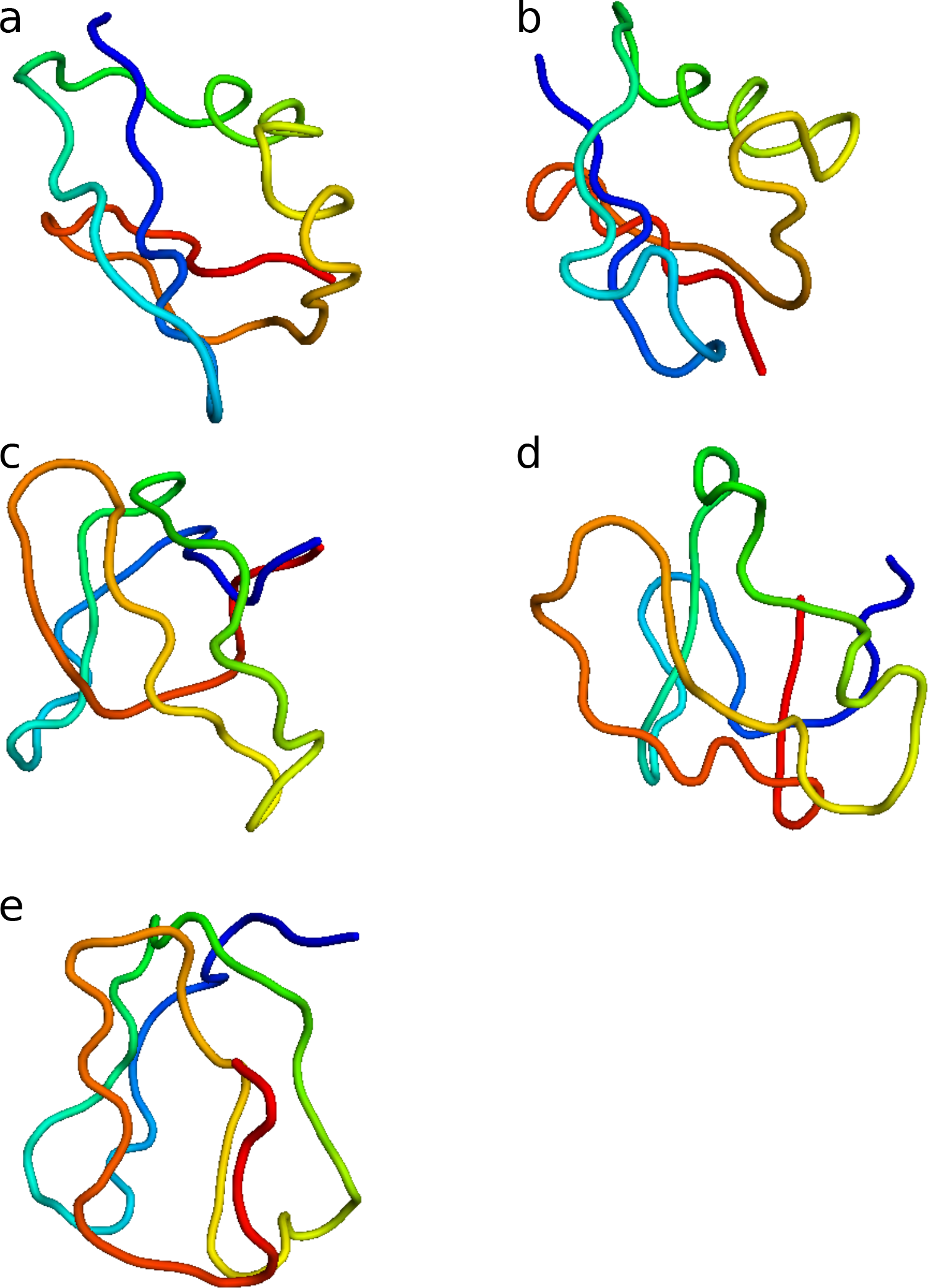
Reconstruction of protein structures from simulated contact frequency data. Shown is the MAP estimate for *n* = 5 states. **(a,b)**: states approximating the conformation of 1PGA. **(c,d)**: state approximating the conformation of 1SHF. **(e)**: “superfluous”, uninformative state.

**Figure S4:**
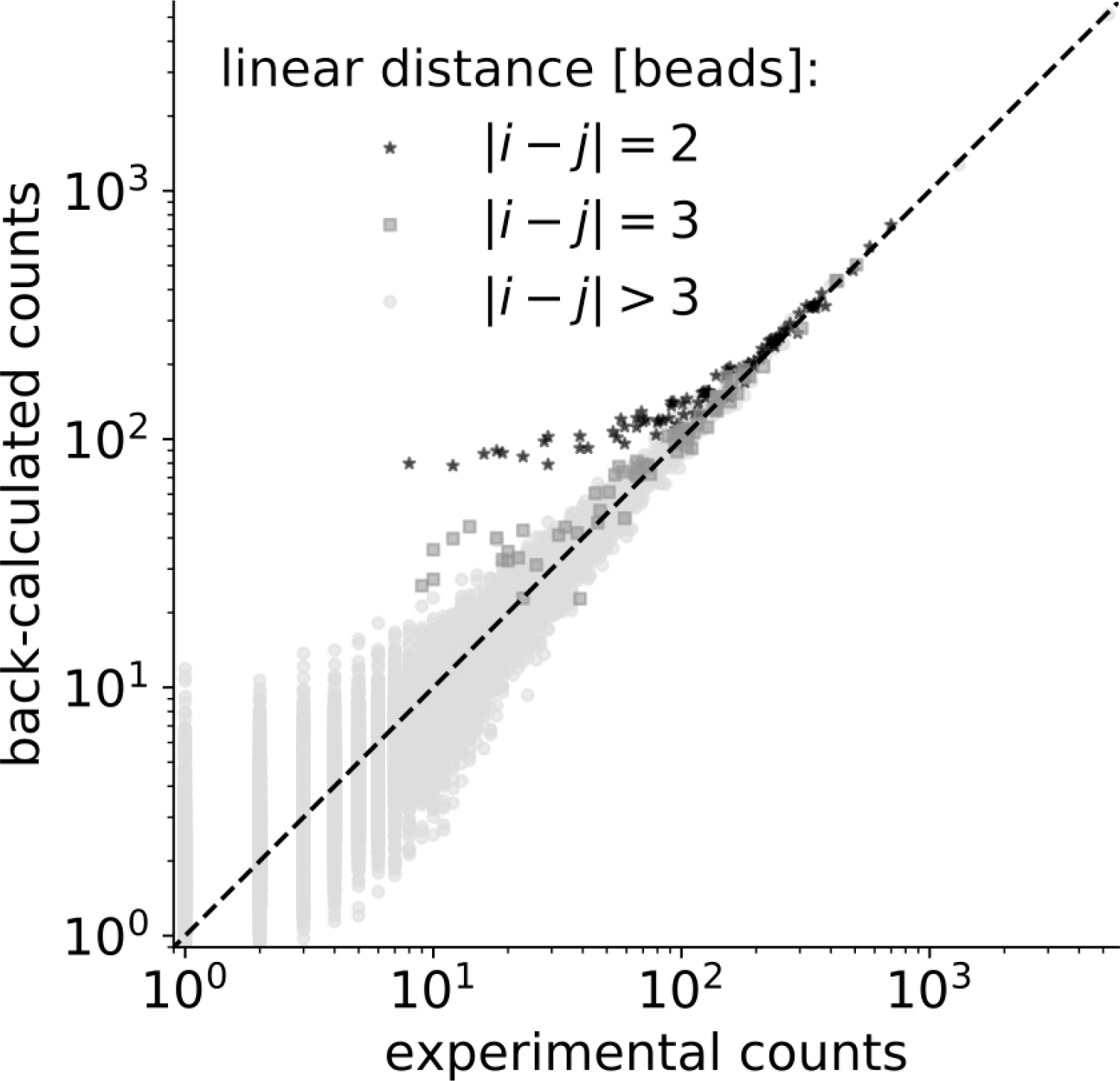
Scatter plot of back-calculated contact counts against experimental counts (input data).

**Figure S5:**
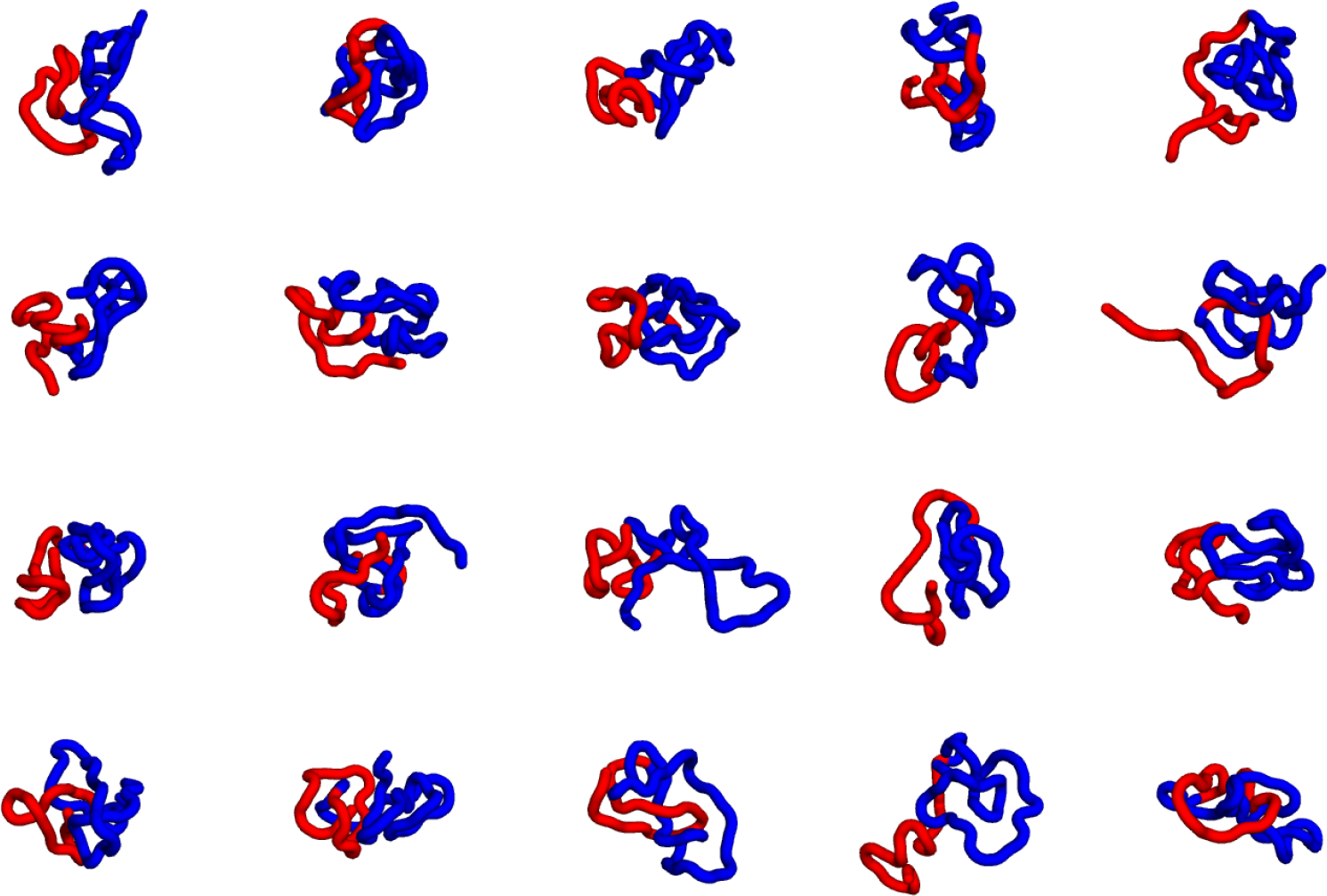
Inference of chromatin structures from 5C data. Shown is the MAP estimate for *n* = 20 states calculated at a low resolution of 15 kb per bead. The *Tsix* and *Xist* TADs are shown in red and blue, respectively.

**Figure S6:**
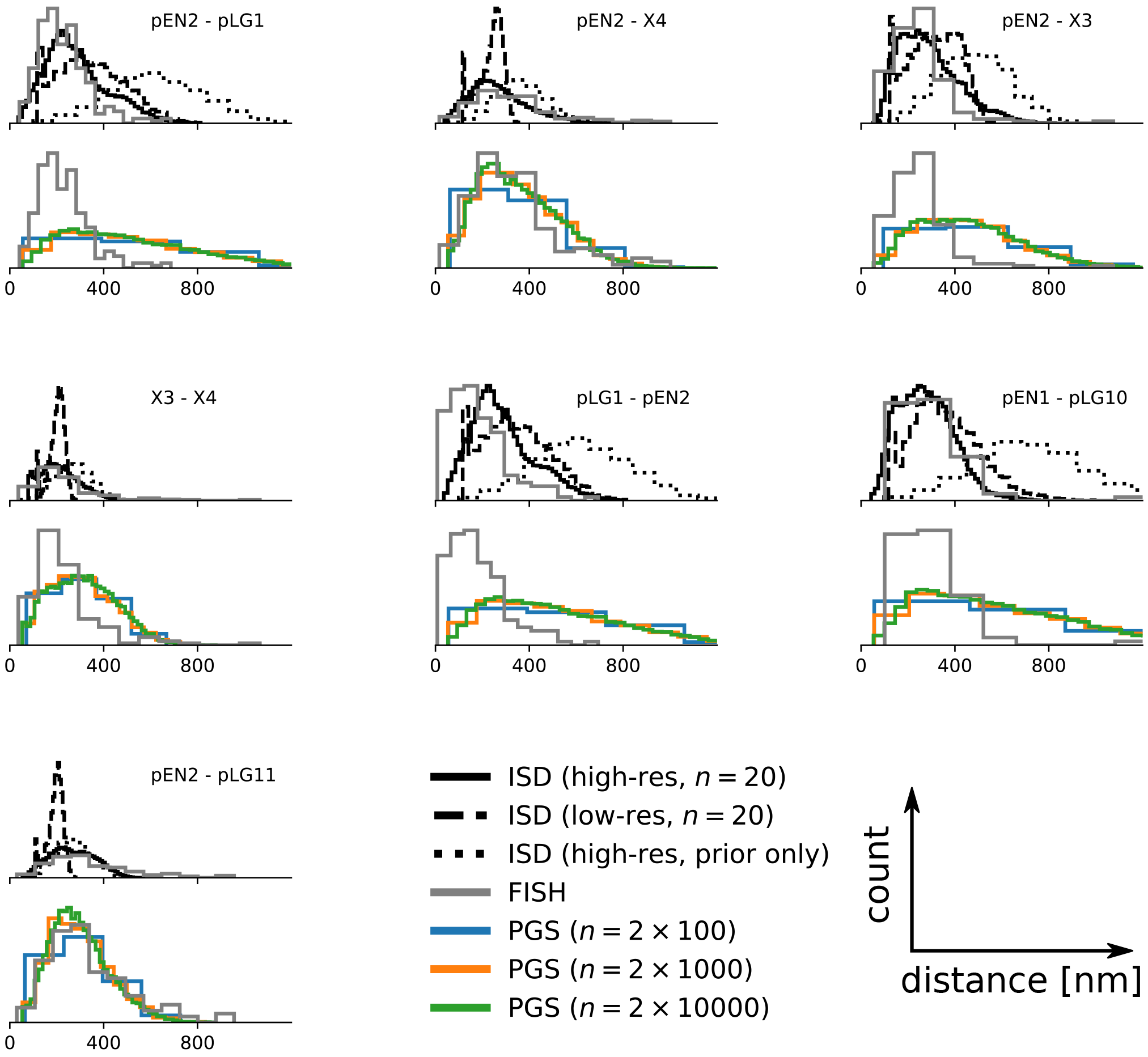
Pairwise distances between several loci as measured by DNA FISH and multi-state models inferred using ISD (top panels) and PGS (bottom panels). Names of genes overlapping the probes are shown.

### Inferential Structure Determination

The Inferential Structure Determination (ISD) approach (24) views biomolecular structure determination as a problem of statistical inference. ISD addresses the following question: “what do we know about the structure of a biological macromolecule given the experimental and prior information?” by means of probable inference. Formally, the answer is given by *p*(*x|D, I*), namely, the conditional probability distribution of the structure *x* conditioned on the data *D* and the prior information *I*. By virtue of Bayes’ theorem, this conditional probability, the so-called *posterior*, can be decomposed as follows:

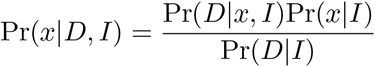

Pr(*D|x, I*) is a statistical model for the data; to emphasize that for given data *D* this probability can be viewed as a function of the structure, we introduce the notation *L*(*x*) = Pr(*D|x, I*) and call *L*(*x*) the *likelihood (function)*. Pr(*x|I*) describes the prior knowledge about the structure and is therefore referred to as the *prior*. The normalization constant Pr(*D|I*) is called the *evidence* and is the average of the likelihood function under the prior, i.e. Pr(*D|I*) = ∫ *L*(*x*) Pr(*x|I*) *dx*.

In general, the likelihood will depend on additional, unknown parameters *θ* that are not of primary interest and therefore called *nuisance parameters*. To infer both the structure and the nuisance parameters from the data, we use their joint posterior probability

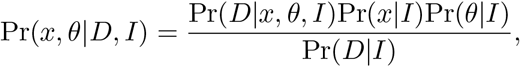

where we assume that the prior distributions of the structure and the nuisance parameters are independent.

In our application of ISD to population Hi-C data, we infer a population of *n* conformers and thus have to replace the single structure *x* in the above equations with a collection *X* of single structures; *X* = (*x*^1^, …, *x*^*n*^). Assuming that the conformers do not interact with each other, the prior distribution Pr(*x*^1^, …, *x*^*n*^) factorizes and the posterior distribution can be written as

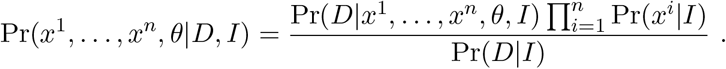

### Statistical model for 5C data

In our model for population contact data, we represent the system by *M* equal-sized beads arranged in a chain. In case of the protein data, a single bead represents one amino acid. In case of the 5C data, a single bead represents 3kb of chromatin similar to the model by Giorgetti *et al.* (20). The mapping between beads and restriction fragments is based on the sequence positions of the restriction sites. If multiple restriction fragments are contained within one bead, all contributing 5C counts are summed and assigned to that bead. If, on the other hand, a restriction fragment spans several, say *p*, beads, all 5C counts involving that restriction fragment are assigned to a central, i.e. the 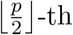, bead. This coarse-graining procedure yields a symmetric contact frequency matrix *D*_*ij*_ with *i, j* ∈ {1, …, *M* }. In our structure calculations, we ignore sequential contacts (|*i − j*| = 1), because they do not contribute any new information which is not already encoded in the structural prior distribution described later. We also neglect data points with zero counts. A script implementing these steps starting from the raw 5C data (5) is provided with the Python code implementing our approach.

In Bayesian data analysis, the likelihood quantifies how well a model matches the data. It is convenient to decompose the likelihood into a *forward model f*, which back-calculates data 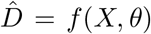 from the structure population *X* and the nuisance parameters *θ*, and an *error model g*, which maps the mock data 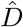 to the probability 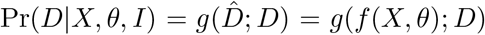. The error model accounts for both experimental noise and the error inherent in the approximate forward model.

We first describe a forward model which back-calculates a contact frequency matrix 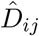 from the set of *n* states *X* = (*x*^1^, …, *x*^*n*^). For each state *x*^*k*^, we calculate a contact matrix with entries 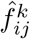 such that

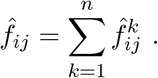

If we define a distance *d*_*c*_ below which two beads are defined to be in contact, ideally, 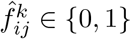 is a binary-valued quantity. But because we use a gradient-based MCMC sampling algorithm, we require all occuring probabilities to be differentiable and thus approximate a hard contact between two beads *i* and *j* with a flipped sigmoidal function *s*(*d*) = (*γd*(1 + *γ*^2^*d*^2^)^−1/2^ + 1)/2. Here, *γ* is a parameter determining the steepness of the smoothed contact such that for large *γ* the hard contact criterion is recovered: lim_*γ*→∞_ *s*(*d*) = 1 − θ(*d*).

The total number of contacts between two beads,

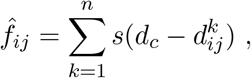

where 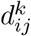 is the distance between beads *i* and *j* in the *k*-th population member. 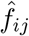 is not directly comparable to the experimental Hi-C interaction frequency matrix *f*_*ij*_, because the number *n* of states does not match the (unknown) number of molecules analyzed in the experiment. To account for this mismatch, we multiply the entries of the back-calculated contact matrix with a positive scaling factor *α* and treat it as a nuisance parameter. The complete forward model then becomes

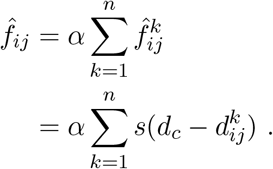

We model the discrepancy between experimental and back-calculated data using a Poisson distribution with rate 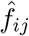:

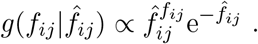

For a single contact, the likelihood is then given by

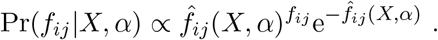

Assuming that the single measurements are independent, the likelihood for the complete data set simply is the product of the likelihood for each individual data point:

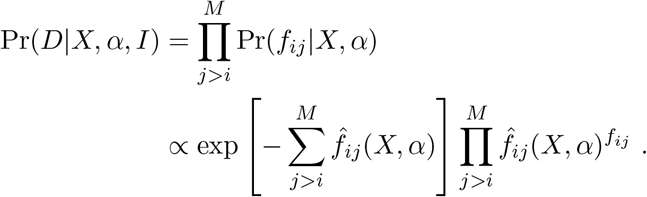

It is important to note that we could have chosen a different error model. In principle, Bayesian model comparison allows us to determine the error model most compatible with the data from a set of choices. However, we restrict this study to a Poisson error model, because the Poisson distribution is discrete and has been used previously to model Hi-C data.

### Statistical models for prior information

#### Conformational prior distribution

We represent chromatin by a beads-on-a-string model composed of *M* beads with radii *r*_*i*_ already presented in our previous work (23, Figure S7). The distance *d*_*ij*_ between the centers of two neighboring beads *i, j* with |*i* − *j*| = 1 is restrained with a harmonic potential *E*_bb_ to values smaller than *r*_*i*_ + *r*_*j*_, that is,

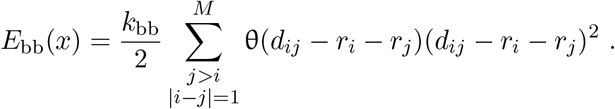

**Figure S7:**
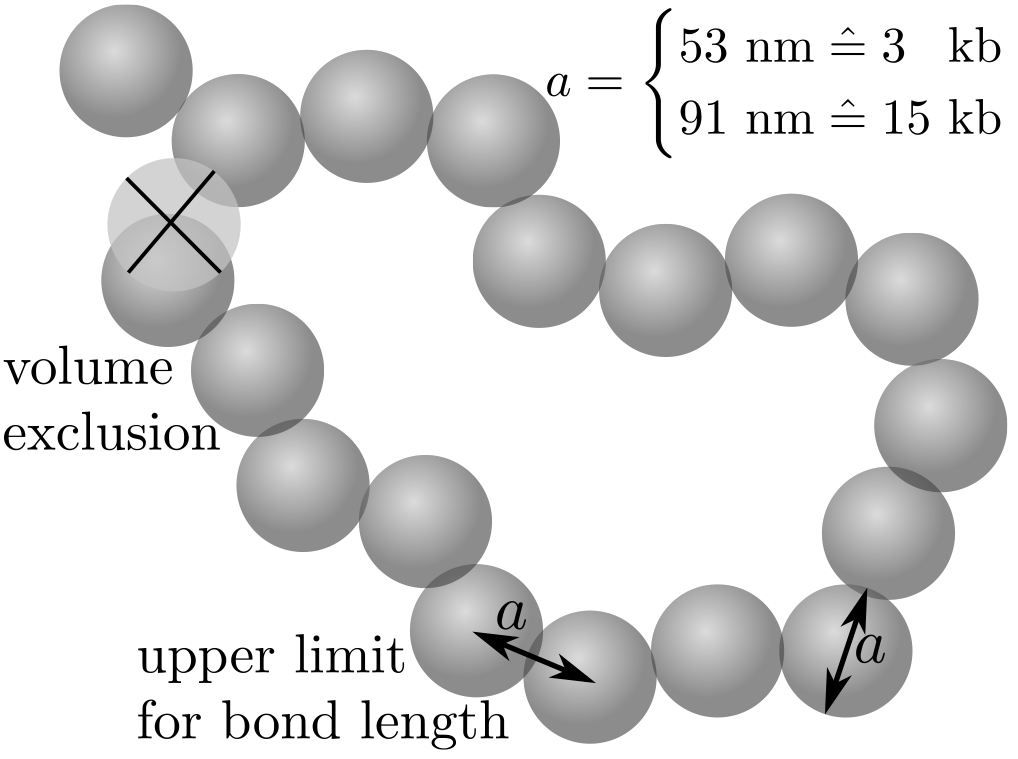
Illustration of the coarse-grained beads-on-a-string model used in this work. Volume exclusion is mediated by a quartic potential, while exceeding the maximum bond length is penalized quadratically.

If any two beads *i, j* are closer than *r*_*i*_ +*r*_*j*_, they experience a repulsive force stemming from a potential *E*_ve_ which scales with 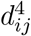;

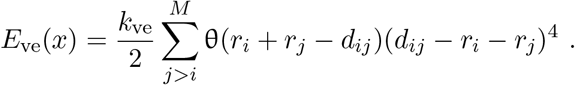

*E*_ve_ serves to emulate a volume exclusion (or self-avoidance) of the chromatin fiber.

As the conformational prior distribution for a single chromosome model we then assume a Boltzmann distribution at inverse temperature *β* = 1, in which the system energy is composed of the backbone and the volume exclusion potential;

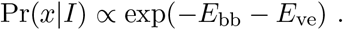

Because we assume that single copies of a chromosome do not interact, the conformational prior distributions for single conformers are independent and we can thus write the conformational prior distribution for the model conformers as

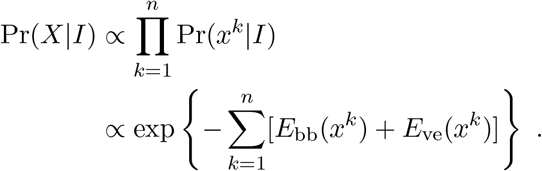

Additionally, when infering protein structures from simulated contact data, we enclose all beads in a spherical, soft shell with radius *r*_sphere_ = 2*r̄ M*^−3^ centered around the coordinate origin. Here, *r̄* is the average bead radius. Beyond this distance from the origin, beads experience a force pushing them back into the spherical shell. The potential of this force acting on a bead *k* is

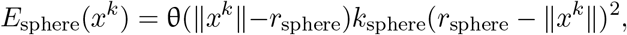

where we set *k*_sphere_ = 5.

#### Prior distribution for *α*

For the Gibbs sampler described in Supplementary Note “Markov Chain Monte Carlo (MCMC) methods for sampling the posterior distribution”, we need to sample from the conditional posterior distributions of both *X* and *α*. Assuming a uniform prior distribution over the positive real numbers, the conditional posterior distribution of the ensemble scaling factor *α* is a Gamma distribution Γ(*ν, β*) with shape *ν* = 1 + ∑_*ij*_ *f̂*_*ij*_ and rate 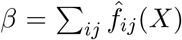. In this work, instead of a uniform prior distribution for *γ*, we use a Gamma distribution with shape *ν*^*P*^ and rate *β*^*P*^. This is a convenient choice, because Γ(*ν, β*)Γ(*ν′*, *β′*) = Γ(*ν* + *ν′* − 1, *β* + *β′*). The conditional posterior distribution for *α* thus becomes

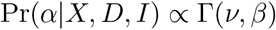

with

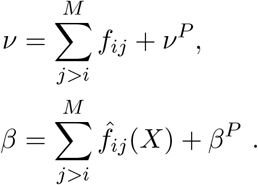

Throughout this work, we set *ν*^*P*^ to half the average of all non-zero contact counts and *β*^*P*^ = 1/*n*.

### Markov Chain Monte Carlo (MCMC) methods for sampling the ISD posterior distribution

Because the posterior distribution Pr(*X, α|D, n, I*) is of non-standard form, we employ a range of MCMC methods to approximate Pr(*X, α|D, n, I*) by a large number of samples. This allows us to define a “structural error bar” and to calculate averages, as opposed to minimization of the posterior, which only yields a maximum *a posteriori* (MAP) estimate with no measure of uncertainty.

#### Gibbs sampling

Using Gibbs sampling (32), we can split up the formidable task of sampling from the joint posterior distribution for the structural model population *X* and the ensemble scaling factor *α* into sampling from the respective conditional distributions. Gibbs sampling is an MCMC algorithm which suceeds by iteratively drawing a sample from the conditional distribution for the first variable, then conditioning the conditional posterior distribution of the second variable on the just drawn value for the first variable and so on, until a sample for each variable has been drawn. These samples are then stored and the above procedure iterated until the collection of all stored samples reasonably approximates the joint distribution. More formally, in our case,

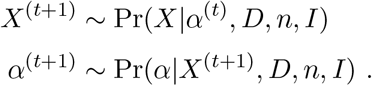

This allows us to draw samples from the conditional posterior distributions for *X* and *α* using appropriate samplers.

#### Sampling from the structural posterior distribution

The most challenging and computationally expensive part in an ISD calculation is drawing representative samples from the conditional posterior distribution for the molecule structure. In our setting, the problem is even harder, because the posterior distribution is not only over one, but several structures. For *n* conformers with *M* beads each, the space on which this probability distribution lives, is 3 × *n* × *M*-dimensional. The 3*M* coordinates of a single conformer are strongly correlated because of bonded and non-bonded interactions encoded in the force field and contact restraints. Because the contact restraints are ensemble restraints, they induce also a strong correlation between coordinates of different conformers.

We approach this difficult sampling problem using Hybrid Monte Carlo (33), a Markov Chain Monte Carlo method which uses the gradient of the log-probability of the probability distribution of interest to guide a short molecular dynamics (MD) trajectory. The endpoints of these trajectories are then used as distant proposal states with high acceptance probability.

Hybrid Monte Carlo requires three parameters, namely, the number of integration steps to calculate the MD trajectory, the integration time step and the mass matrix. Rules-of-thumb to set these parameters are proposed in the literature (34), but require prior knowledge about the probability distribution. We thus use heuristics; setting the mass matrix to unity, the number of integration steps to 100 and the timestep such that on average, ~ 50% of HMC moves are accepted. Strictly speaking, the latter violates detailed balance, but we observe that after a burn-in period, the integration timesteps change very little and the error introduced thus will be negligible.

#### Sampling the ensemble scaling factor *α*

The conditional posterior distribution Pr(*α|X, D, n, I*) is a Gamma distribution (see *Supplementary Note: Statistical models for 5C data and prior information*), for which random number generators are readily available in many programming languages. It is important to note that the Gamma distribution as the conditional distribution for *α* is specific to the Poisson error model and prior we use in this work. For a different choice of error model and / or prior, direct sampling of *α* might not be possible and MCMC methods may have to be employed.

#### Replica Exchange

Because of multimodality, the described Gibbs sampler can get trapped in high-probabiltiy regions and sampling thus becomes inefficient in the best and incorrect in the worst case. To alleviate this issue, we embed the Gibbs sampler in a Replica Exchange (35) algorithm. Replica Exchange simulates not only the probability distribution of interest, but a family of distributions parameterized by one or several temperature-like parameters. At high temperatures, sampling is more likely to be exhaustive and by occasionally attempting to swap configurations between neighbouring temperatures, configurations from the high-temperature simulations can diffuse to the low-temperature simulations. RE thus facilitates sampling by increasing hopping between modes in low-temperature simulations.

In this work, we introduce a single replica parameter *β*, analogous to the inverse temperature in a Boltzmann ensemble. This parameter changes the incluence of the likelihood function and the volume exclusion, while the connectivity restrained encoded in *E*_bb_ remains fully switched on. The family of tempered posterior distributions we simulate then is

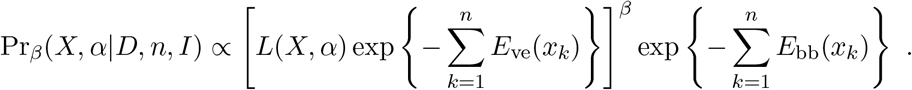

In our simulations, *β* assumes values in the interval [10^−6^, 1]. Over the course of several simulations, the sequence of *β* values is optimized.

